# A unified transcriptome dataset for Amaryllidoideae species

**DOI:** 10.1101/2025.11.24.690262

**Authors:** Karen Cristine Gonçalves dos Santos, Natacha Merindol, Isabel Desgagné-Penix

**Affiliations:** Département de Biochimie, Chimie, Physique, et Science Forensique, Université du Québec à Trois-Rivières, Trois-Rivières, QC, Canada; Plant Biology Research Group, Université du Québec à Trois-Rivières, Trois-Rivières, QC, Canada

## Abstract

Amaryllidoideae plants produce structurally diverse and unique alkaloids with potent anti-cholinesterase, antiviral, and antitumor activities, making this subfamily a rich source of pharmaceutical leads. Despite the absence of reference genomes for any Amaryllidoideae species, many enzyme characterization and pathway reconstruction efforts to date have been made possible through transcriptome mining, often requiring bioinformatic expertise and data preprocessing. To facilitate new studies in this subfamily, here we present AmarylOmicBase, a unified transcriptomic dataset that integrates assemblies, annotations, and expression profiles from 39 studies, covering 27 species and four hybrid cultivars across 13 genera of Amaryllidoideae. The AmarylOmicBase includes both published and *de novo* assemblies generated from published raw data using Trinity or IsoSeq workflows and provides standardized functional annotation and quantitative expression datasets. AmarylOmicBase provides ready-to-use datasets that support gene discovery, comparative transcriptomics, and pathway-level investigations for specialized metabolism, including Amaryllidaceae alkaloid biosynthesis. By providing ready-to-use datasets and fully reproducible analysis scripts, this resource reduces computational barriers and expands access to transcriptomic information for researchers working on non-model plant species. AmarylOmicBase provides a centralized resource for transcriptomic data that can be reused in studies of enzyme function, pathway evolution, and regulatory processes in Amaryllidoideae.

## Background & Summary

The Amaryllidoideae subfamily, of the Amaryllidaceae plant family, comprises approximately 60 genera and more than 800 species distributed worldwide. Members of this subfamily have attracted sustained research interest due to their distinctive specialized metabolism, particularly the production of structurally diverse alkaloids with reported potent pharmaceutical activities including anti-cholinesterase (galanthamine ^[1–3]^), antiviral (lycorine, haemanthamine, and pancracine, among others ^[4–6]^), and antitumor (lycorine, narciclasine, and haemantamine, among others ^[7,8]^) properties. This chemical diversity has motivated numerous biochemical, molecular, and evolutionary studies aimed at understanding enzyme function, pathway organization, and the genetic basis of metabolite diversification in Amaryllidoideae species.

To date, no reference genome is available for any Amaryllidoideae species, largely due to large genome sizes and complex genome architectures characteristic of this family. This has severely hindered progress in elucidating the pathways of these pharmaceutically relevant compounds for heterologous production. In this context, transcriptome sequencing has become the primary source of gene sequence information and has enabled many studies focused on enzyme characterization, pathway reconstruction, and comparative analyses. For instance, the first transcriptome assemblies of this subfamily led to the identification and characterization of several enzymes involved in these pathways such as norbelladine 4’-*O*-methyltransferase (N4OMT), noroxomaritidine/norcraugsodine reductase (NR) and the phenol-coupling cytochrome P450 96T (CYP96T) in *Narcissus aff*. *pseudonarcissus*, *Galanthus elwesii* and *Galanthus* sp. ^[9–11]^. Later, the assembly of *N. pseudonarcissus* King Alfred’s transcriptome led to the identification and characterization of norbelladine synthase (NBS^[12]^). Transcriptome mining of *Lycoris aurea* ^[13]^ and *Leucojum aestivum* ^[14]^ allowed the characterization of two cytochrome P450s in the phenylpropanoid pathway, namely cinnamate 4-hydroxylase (C4H or CYP73A) and *p*-coumaroyl 3′-hydroxylase (C3’H or CYP98A) ^[15,16]^. Finally, two recent studies successfully reconstituted the Amaryllidaceae alkaloid network in *Nicotiana benthamiana* through transcriptome mining of *L. aestivum* ^[17]^ and *Narcissus* cv. ‘Tête-à-Tête’ ^[18]^, further demonstrating the power of transcriptomic data for pathway elucidation. Over the past decade, RNA sequencing datasets have been generated for multiple Amaryllidoideae species, tissues, and experimental conditions and deposited across public repositories.

Despite these advances, transcriptome-based discovery remains technically challenging. Datasets are dispersed across repositories, generated using different sequencing platforms and assembly strategies, and vary in annotation quality and expression quantification, resulting in heterogeneous data formats and variable quality. This fragmentation creates barriers to reproducibility, comparative analysis, and systematic exploration of enzyme diversity — particularly relevant given that the alkaloid profiles differ markedly among Amaryllidoideae species ^[19]^. In addition, raw data processing, functional annotation, and expression quantification typically require bioinformatic expertise and access to computational infrastructure, which can limit accessibility for researchers working on non-model plant species.

To facilitate data reuse and comparative analyses within the Amaryllidoideae, we developed AmarylOmicBase, a unified transcriptomic dataset that integrates publicly available RNA-seq datasets from this subfamily into a standardized and reproducible framework. AmarylOmicBase compiles transcriptome assemblies, functional annotations, and quantitative expression profiles from 39 independent studies, encompassing 27 species and four hybrid cultivars across 13 genera. The dataset includes de novo assemblies generated from publicly available raw sequencing data using consistent short-read (Trinity) and long-read (Iso-Seq) workflows and were then processed through standardized pipelines for quality assessment, functional annotation, and expression quantification, enabling comparison across species, tissues, and experimental conditions. In addition, all analysis scripts and workflows are provided to ensure full reproducibility and to support future updates as new datasets become available.

By consolidating fragmented transcriptomic resources into a single, curated platform, AmarylOmicBase provides ready-to-use datasets that support gene discovery, comparative transcriptomics, and pathway-level investigations in Amaryllidoideae. This resource lowers technical barriers to transcriptomic data reuse and expands access to molecular information for research on specialized metabolism, pathway evolution, and regulatory processes in this non-model plant subfamily.

## Methods

### Selected datasets and assembly pipeline

The dataset was constructed following the pipeline summarized in Figure 1 and Table 1 shows the accessions for the raw data used to generate each assembly^[20–64]^. For SRP403412^[57]^, only the wild type callus samples were used. Raw data were downloaded from the Sequencing Read Archive (SRA) using sra-toolkit 3.0.9, then quality control was performed using fastp 0.23.4 ^[65]^ (mean quality = 20; unqualified percent limit = 30; cut-front window-size = 5; cut-right window-size = 4; cut-right mean quality = 15; length required = 50). Samples from the projects SRP067647, SRP067648, and SRP067700 ^[9,10]^ were sequenced with paired-end reads of 50 bp. Thus, the minimum length after trimming was set to 30 bp. The *Lycoris radiata* assembly from Park, et al. ^[66]^ (SRR8799443) was not included in the dataset because no raw data were available.

**Figure 1.**
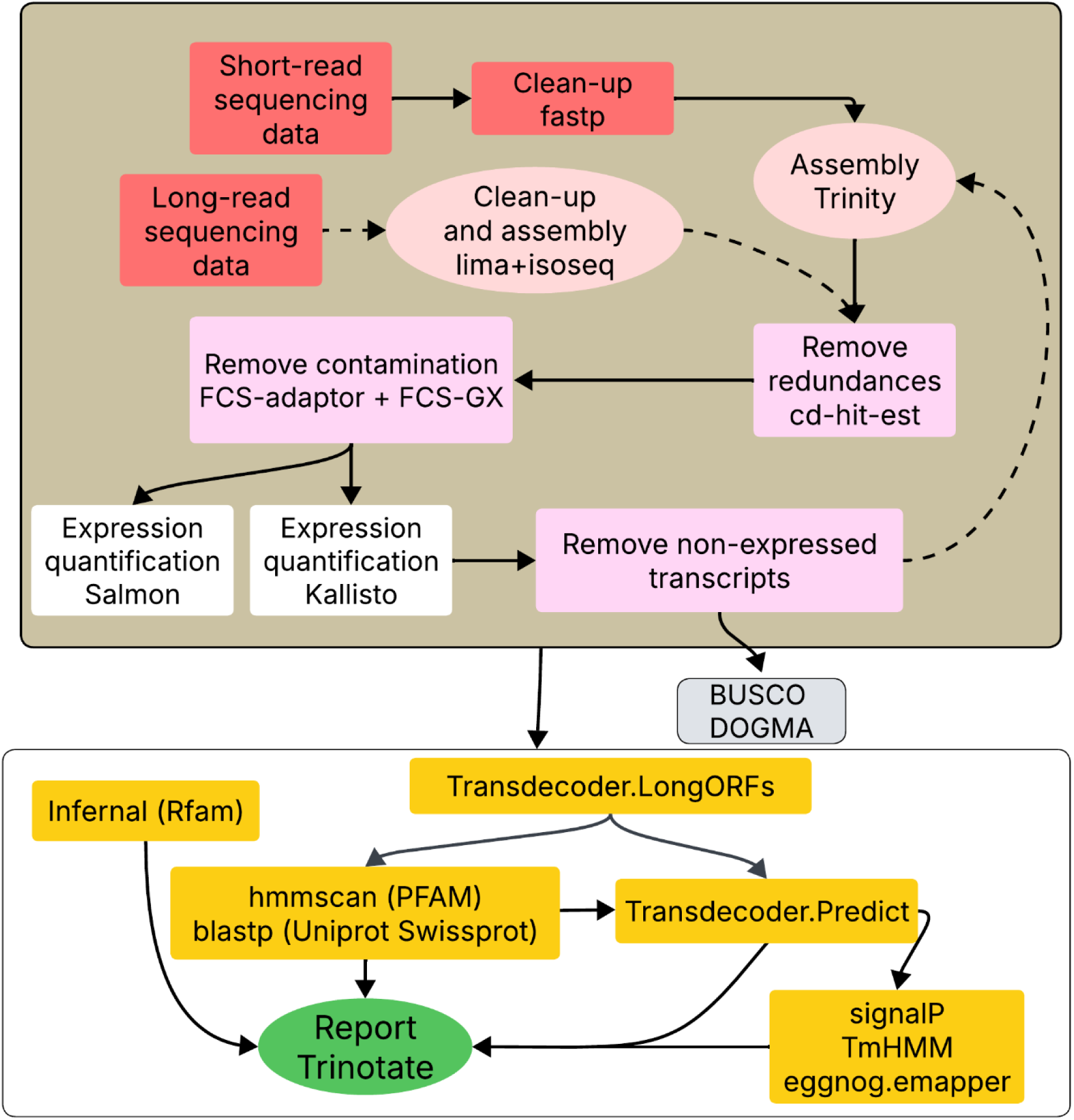
AmarylOmicBase construction pipeline. Short reads were downloaded from the Sequence Read Archive (SRA) then trimmed and filtered with fastp. Reads were then assembled with Trinity and simplified with CD-HIT-EST. When long read data was available, they were downloaded from SRA, preprocessed with lima and clustered into full-length non-chimeric reads with IsoSeq. Assemblies were cleaned up using FCS-adaptor followed by FCS-GX. Reads were pseudoaligned to the assemblies with kallisto. The result was used to filter our assemblies further, removing Trinity isoforms that were not expressed. For species/cultivars with long and short read data, the cleaned up full-length non-chimeric reads were used as input, along with the short reads, for Trinity. This hybrid assembly was processed as previously described. Assemblies were analyzed with BUSCO and DOGMA for estimation of completeness. Transcripts were also analyzed with INFERNAL for the prediction of secondary RNA structure. Additionally, an initial coding sequence identification was performed with Transdecoder.LongORFs and the result was used for homology-based annotation with BLASTp using UniProt SwissProt, and detection of protein family domains, hmmscan using the Pfam-A dataset. Results were entered into TransDecoder.Predict, and the improved open reading frame predictions were used to search signal peptides and transmembrane domains, with SignalP and TmHMM, respectively, and for orthologous group

**Table 1.** Raw data included in the dataset (SRA project ID, individual SRA run IDs are provided when the project included data from multiple species).

| Species | Accession | Sequencing technology | Reference |
| --- | --- | --- | --- |
| *Amaryllis belladonna* | ERP023948<sup>[31]</sup>; ERR2040723 | Illumina HiSeq 2000 | [67] |
| *Clivia miniata* | SRP152913<sup>[27]</sup> | Illumina HiSeq 2500 | [68] |
|  | SRP363321<sup>[41]</sup> | Illumina MiniSeq | [69] |
|  | SRP507161<sup>[58]</sup>; SRR29002427 | DNBSEQ-T7 | [70] |
| *Crinum asiaticum* | SRP507161<sup>[58]</sup>; SRR29002426 | DNBSEQ-T7 | [70] |
|  | SRR29002425 |  |  |
| *Crinum x powellii* | SRP434863<sup>[47]</sup> | Illumina NovaSeq 6000 | [71] |
| *Galanthus elwesii* | SRP067648<sup>[20]</sup> | Illumina HiSeq 2000 | [11] |
| *Galanthus sp.* | SRP067700<sup>[21]</sup> | Illumina HiSeq 2000 | [11] |
| *Hippeastrum striatum* | SRP507161<sup>[58]</sup>; SRR29002422-3 | DNBSEQ-T7 | [70] |
| *Hippeastrum vittatum* | SRP388119<sup>[48]</sup> | Illumina NovaSeq 6000 | [72] |
| *Hippeastrum* hybrid cv. Blossom Peacock | SRP075443<sup>[24]</sup> | Illumina HiSeq 2000 | [73] |
| *Hippeastrum* hybrid cv. Jewel | SRP250867<sup>[42]</sup> | Illumina NextSeq 500 | [74] |
| *Hippeastrum* hybrid cv. Royal Velvet | SRP427234<sup>[49]</sup> | Illumina HiSeq X Ten | [75] |
| *Leucojum aestivum* | SRP314214<sup>[39]</sup> | Illumina HiSeq 2000 | [14] |
| *Lycoris aurea* | SRP266367<sup>[54]</sup> | HiSeq X Ten | [76] |
|  | SRP505073<sup>[60]</sup> | PacBio Sequel | [77] |
|  | SRP505105<sup>[62]</sup> | Illumina HiSeq X Ten |  |
|  | SRP507161<sup>[58]</sup>; SRR29002421 | DNBSEQ-T7 | [70] |
|  | SRP484118<sup>[34]</sup> | Illumina HiSeq X Ten | [78] |
|  | SRP501977<sup>[64]</sup> | Illumina NovaSeq 6000 | [79] |
| *Lycoris chinensis* | SRP266062<sup>[52]</sup> | HiSeq X Ten | [76] |
|  | SRP379054<sup>[44]</sup> | Illumina HiSeq 2500 | [80] |
| *Lycoris incarnata* | SRP267213<sup>[55]</sup> | Illumina HiSeq X Ten | [76] |
| *Lycoris longituba* | SRP161854<sup>[33]</sup> | Illumina HiSeq 4000 | [81] |
|  | SRP230579<sup>[37]</sup> | Illumina HiSeq X Ten | [82] |
|  | SRP313739<sup>[63]</sup> | Illumina NovaSeq 6000 | [83] |
|  | SRP266290<sup>[53]</sup> | Illumina HiSeq X Ten | [76] |
| *Lycoris radiata* | SRP223567<sup>[54]</sup> | BGISeq-500 | [84] |
|  | SRP266051<sup>[50]</sup> | Illumina HiSeq X Ten | [76] |
|  | PRJCA006232<sup>[85]</sup> | Illumina HiSeq 2500 | [86] |
|  | SRP404775<sup>[56]</sup> | DNBSEQ-T7 | [87] |
|  | SRP507161<sup>[58]</sup>; SRR29002417-9 | DNBSEQ-T7 | [70] |
| *Lycoris sprengeri* | SRP310694<sup>[38]</sup> | Illumina HiSeq 2500 | [88] |
|  | SRP266056<sup>[51]</sup> | Illumina HiSeq X Ten | [76] |
| *Narcissus papyraceus* | SRP117744<sup>[28]</sup> | Illumina HiSeq 2000 | [89] |
| *Narcissus pseudonarcissus* | SRP110821<sup>[26]</sup> | Illumina HiSeq 2000 | [12] |
|  | SRP167005<sup>[29]</sup> | Illumina HiSeq 4000 | [90] |
| *Narcissus* cv. 'Tête-à-Tête' | SRP480318<sup>[59]</sup> | PacBio Sequel | [18] |
|  |  | Illumina NovaSeq 6000 |  |
| *Narcissus* sp. aff. *pseudonarcissus* | SRP067647<sup>[23]</sup> | Illumina HiSeq 2000 | [9] |
| *Narcissus tazetta* | SRP083092<sup>[35]</sup> | Illumina HiSeq 2000 | [91] |
|  | SRP107601<sup>[25]</sup> | Illumina HiSeq 2000 | [92] |
|  | SRP158548<sup>[32]</sup> | Illumina HiSeq 2000 | [93] |
|  | SRP385008<sup>[46]</sup> | Illumina NovaSeq 6000 | [94] |
|  | SRP371855<sup>[45]</sup> | Illumina NovaSeq 6000 | [95] |
|  | SRP330736<sup>[40]</sup> |  |  |
|  | SRP403412<sup>[57]</sup> | Illumina HiSeq 2500 | [96] |
|  | SRP186233<sup>[36]</sup> | Illumina HiSeq 2000 | [97] |
| *Narcissus viridiflorus* | ERP023948<sup>[31]</sup>: ERR2040724-5 | Illumina HiSeq 2000 | [67] |
| *Phycella* sp. aff. *cyrtanthoides* | ERP023948<sup>[31]</sup>: ERR3487375 | Illumina HiSeq 2000 | [67] |
| *Rhodophiala pratensis* | ERP023948<sup>[31]</sup>: ERR2040726<sup>[31]</sup> | Illumina HiSeq 2000 | [67] |
| *Scadoxus multiflorus* | SRP507161<sup>[58]</sup>: SRR29002424 | DNBSEQ-T7 | [70] |
| *Traubia modesta* | ERP023948<sup>[31]</sup>: ERR2040727 | Illumina HiSeq 2000 | [67] |
| *Zephyranthes candida* | SRP354596<sup>[43]</sup> | Illumina NovaSeq 6000 | [98] |
|  | SRP507161<sup>[58]</sup>: SRR29002416 | DNBSEQ-T7 | [70] |
| *Zephyranthes carinata* | SRP507161<sup>[58]</sup>:<br>SRR29002415 | DNBSEQ-T7 | [70] |
| *Zephyranthes treatiae* | ERP023948<sup>[31]</sup>: ERR2040728 | Illumina HiSeq 2000 | [67] |

Assemblies were carried out with Trinity 2.14.0 ^[99]^, default parameters. Trinity cannot process sample files containing both paired-end and single-end datasets. Therefore, single-end reads were concatenated with forward reads into a single file, and the script was modified accordingly. Next, assemblies were simplified by removing redundant transcripts with CD-HIT-EST 4.8.1 (

In the case of *Lycoris aurea* and *Narcissus* cv. ‘Tête-à-Tête’, long-read datasets were available. Thus, these datasets were obtained in their original BAM format from SRA, cleaned up with lima, and assembled using isoseq from smrtlink-sequel2 13.1.0.221970 (default settings). The short-read datasets were assembled using Trinity, with the full-length, non-chimeric reads from Iso-Seq in the option. Both isoseq (PB) and Trinity-hybrid (TH) assemblies were then processed with CD-HIT-EST as described above.

Assemblies were analyzed to remove contaminant sequences (from viruses, archeobacteria, bacteria, fungi or animals, as well as adaptors) using FCS-adaptor version 0.5.0 and FCS-GX version 0.5.5^[100]^ on Galaxy Project^[100]^. Expression quantification was performed using Kallisto version 0.46.1^[101]^ and Salmon version 1.4.0^[102]^, with the support scripts align_and_estimate_abundance.pl and abundance_estimates_to_matrix.pl from Trinity, with default settings for both. Trinity isoforms not expressed in any sample (total read count = 0), based on Kallisto’s results, were not annotated.

The assembly quality was assessed with BUSCO 5.7.0^[103]^, using the liliopsida_odb10 dataset. Additionally, transcriptomes were analyzed with DOGMA^[104]^ 1.0.2, based on the analyses from RADIANT 1.1.5, using the database radiant_db_pfam37.3 and the monocots reference set. Assemblies generated by the One Thousand Plant Transcriptomes Initiative ^[67]^ were used to compare completeness against the ones generated in the current study. They were downloaded from the GigaDB^[105]^ using assembly accession codes: LDME for *Amaryllis belladonna*, TRRQ for *Narcissus viridiflorus*, DMIN for *Phycella aff. cyrtanthoides*, JDTY for *Rhodophiala pratensis*, ZKPF for *Traubia modesta*, and DPFW for *Zephyranthes treatiae*.

### Annotation pipeline

Next, the Trinotate 4.0.0 pipeline (http://trinotate.github.io/)^[106]^ was used, with parameters for each software set following Trinotate suggestions. Coding sequences were predicted with TransDecoder 5.5.0 (http://transdecoder.github.io/). This initial prediction of coding sequences was annotated with BLASTp (against Uniprot SwissProt dataset release 2024_04; blast+ 2.13 ^[107]^, evalue=10^-5^, maximum number of targets=1), hmmscan (against Pfam-A dataset version 37.3; HMMER 3.2.2 ^[108,109]^, default settings). Afterward, the outputs of these two analyses were used to improve coding sequence predictions with TransDecoder.Predict, using the option “-- single_best_only”. This final prediction was used for annotation with eggnog-mapper 2.1.12 ^[110]^ and prediction of signal peptides as well as transmembrane domains (using SignalP6 ^[111,112]^ and TmHMM 2.0c ^[113]^, respectively), with default settings for all of them. The transcriptome assemblies were also analyzed with Infernal 1.1.4 ^[114]^ for identification of sequences with secondary RNA structure, using Trinotate’s “—run ‘infernal’” command. These results were compiled into a report using the Trinotate report.

### Data Records

We deposited in Zenodo 14 compressed files ^[115]^ that contain transcriptome assemblies, predicted protein sequences, expression quantification, metadata of samples included in the assemblies, and annotation results for the 31 species and hybrid cultivars included in this study (Table 2). Inside each compressed archive, files are named by species (eg. Lycoris_incarnata.fa).

**Table 2.**
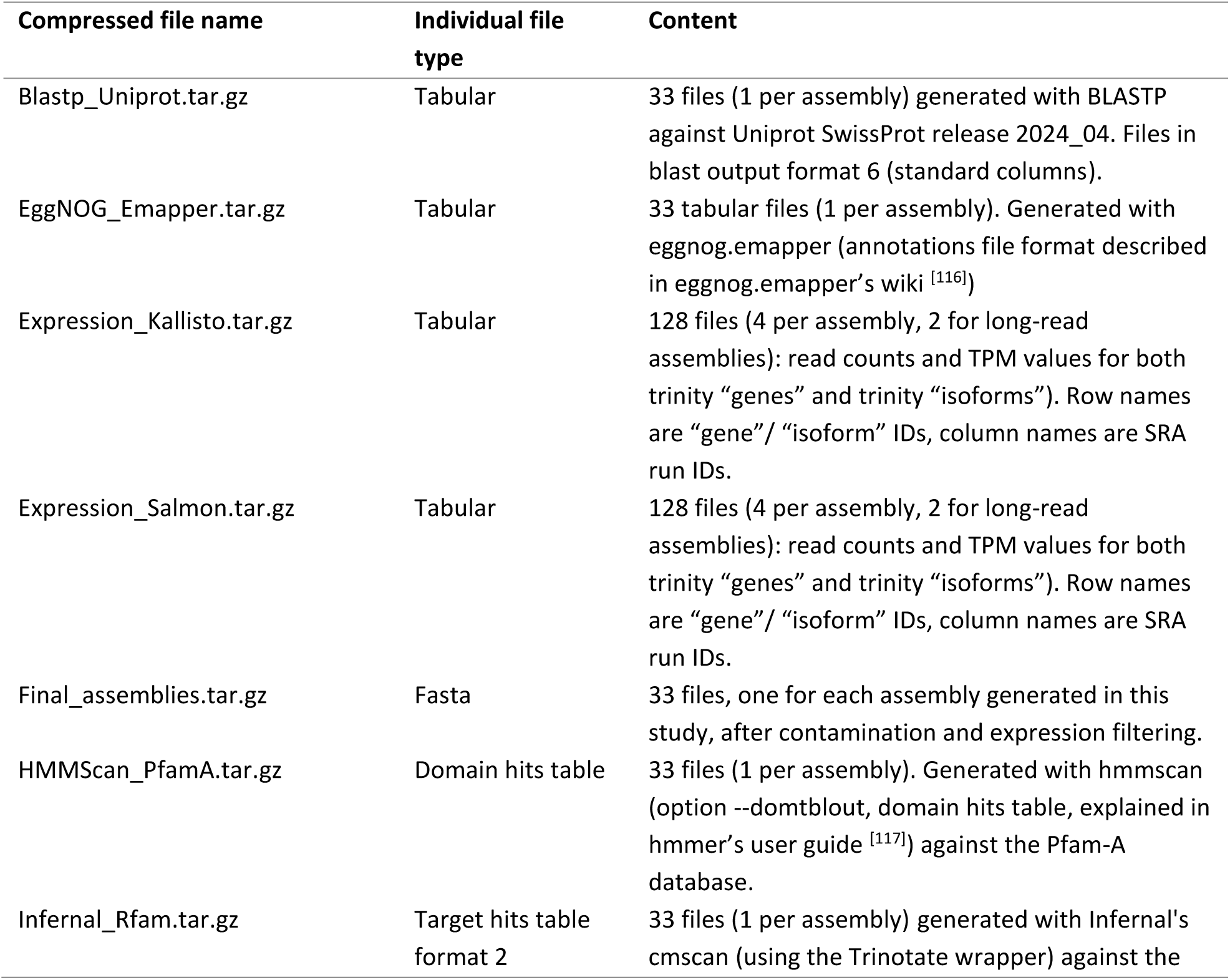
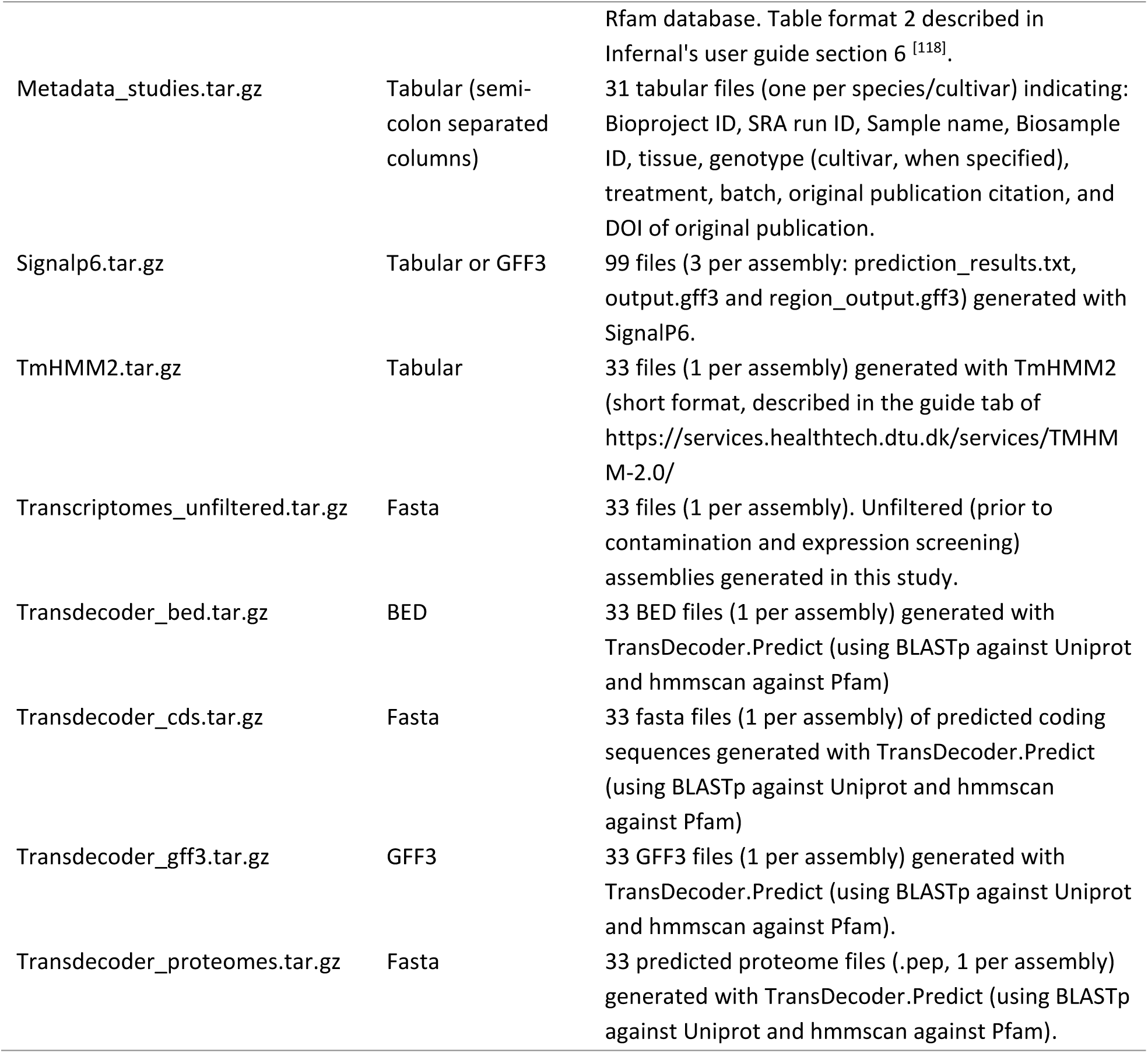
Raw data included in the dataset. Accessions are provided as SRA project ID, with individual SRA run IDs if the project contained data for multiple species).

### Data overview

Most assemblies constructed using Trinity generated a high number of gene and transcripts (Figure 2), with the biggest assemblies (in terms of number of transcripts) being that of *Lycoris radiata*, with 1,155,842 trinity “genes” and 1,690,485 trinity “transcripts”. The smallest assembly was that of *A. belladonna*, with only 30,999 trinity “genes” and 41,679 trinity “transcripts”. Short-read assemblies had lower proportions of “genes” predicted to encode proteins (Figure 3). While 92% and 81.6% of *L. aurea* (PB) and *Narcissus* cv. ‘Tête-à-Tête’ (PB) transcripts, respectively, contained coding sequences, this number dropped to 54.3% for *N. aff. pseudonarcissus* and 42.7% for *A. belladonna*. The assembly with the lowest proportion of protein-encoding genes was *Hippeastrum striatum*, with only 9.6%.

**Figure 2.**
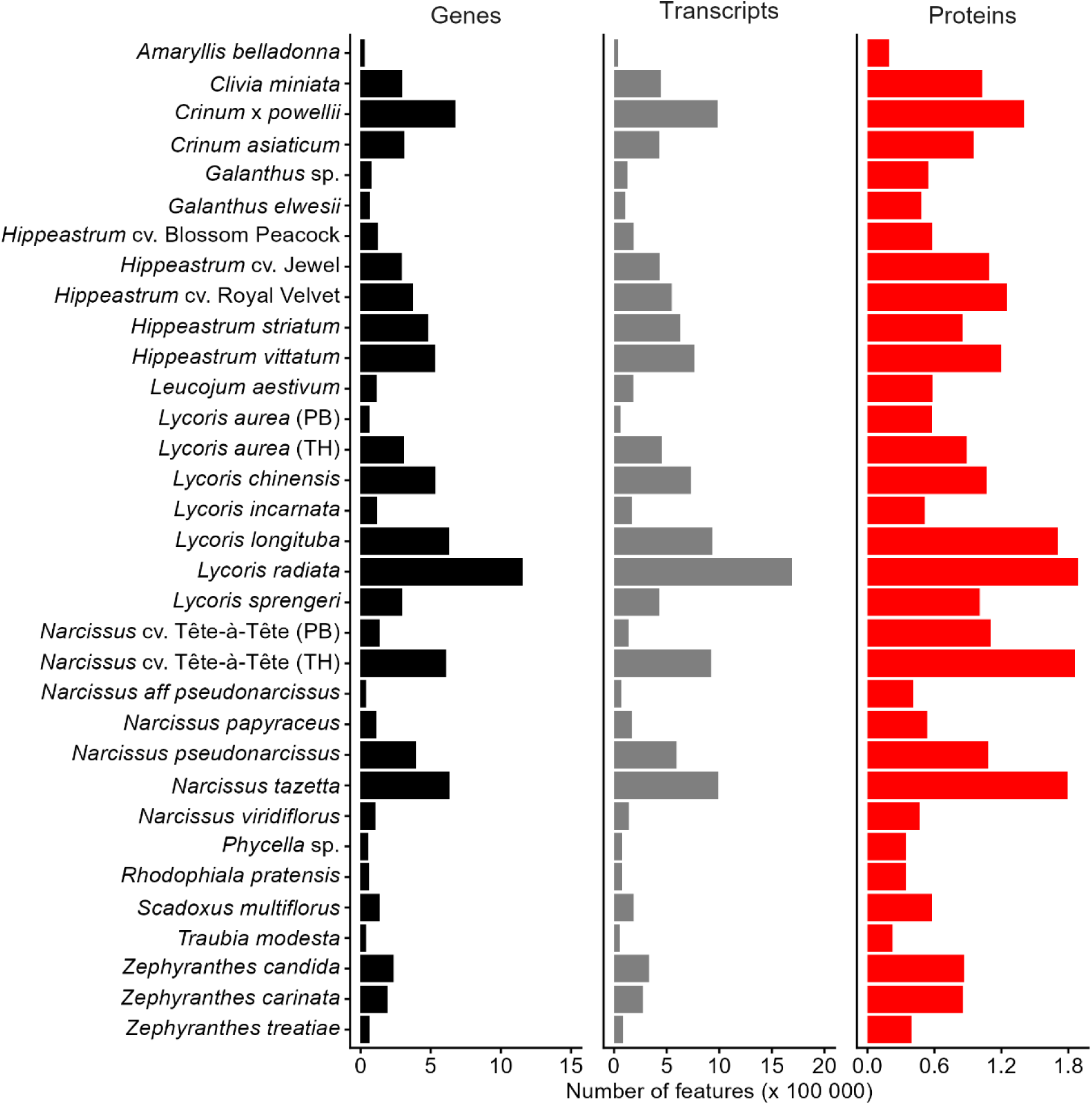
Number of genes, gene isoforms (transcripts), and predicted proteins in each assembly. For *Lycoris aurea* and *Narcissus* cv. ‘Tête-à-Tête’, two assemblies were generated, one using only long-read data (PB) and another using both long- and short read data with Trinity (TH).

**Figure 3.**
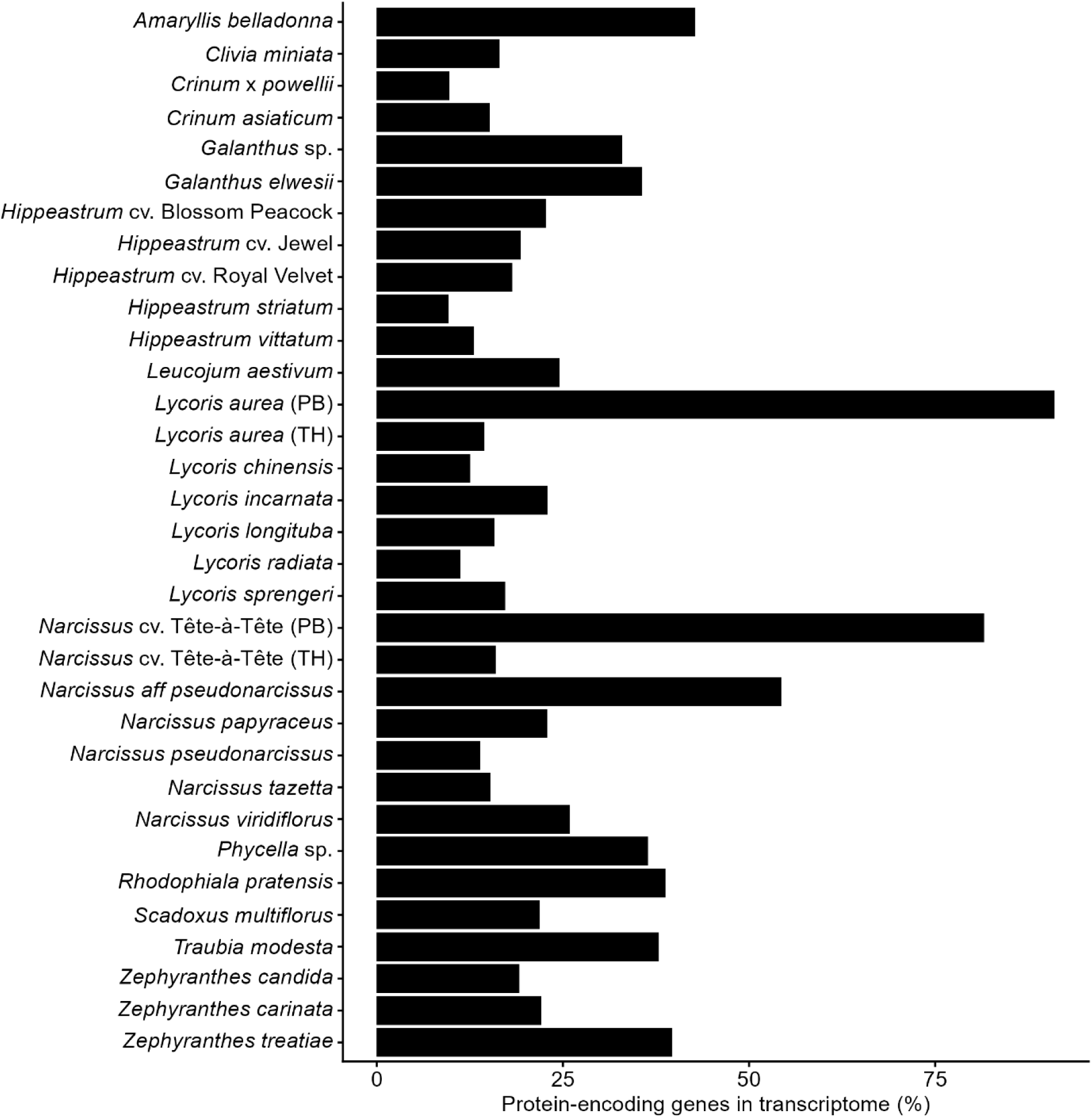
Proportion of protein-encoding genes (in relation to the total number of “genes”) in each transcriptome assembly. For *Lycoris aurea* and *Narcissus* cv. ‘Tête-à-Tête’, two assemblies were generated, one using only long-read data (PB) and another using both long- and short read data with Trinity (TH).

The two assemblies generated with long reads showed the highest proportion of genes annotated using any of the protein-based datasets (Figure 4A) The most annotated assembly was *L. aurea* (PB), with 90% of the transcripts annotated, followed by *N.* cv. ‘Tête-à-Tête’ (PB) with 77.9% of its transcripts annotated. Nevertheless, when considering only ‘genes’ with a predicted coding sequence, all assemblies had most of their ‘genes annotated (Figure 4B). For reference, assembly of the *H. vittatum* transcriptome had the lowest proportion of protein-coding ‘genes’ annotated, at 72.4%. Assemblies were also annotated with Infernal for a homology-based search against the Rfam dataset (Figure 4C), which includes non-coding and cis-regulatory RNA. The proportion of ‘genes’ annotated with Infernal in each assembly was low, ranging between 0.09% for *Z. treatiae* and 0.91% for *L. aurea* (PB).

**Figure 4.**
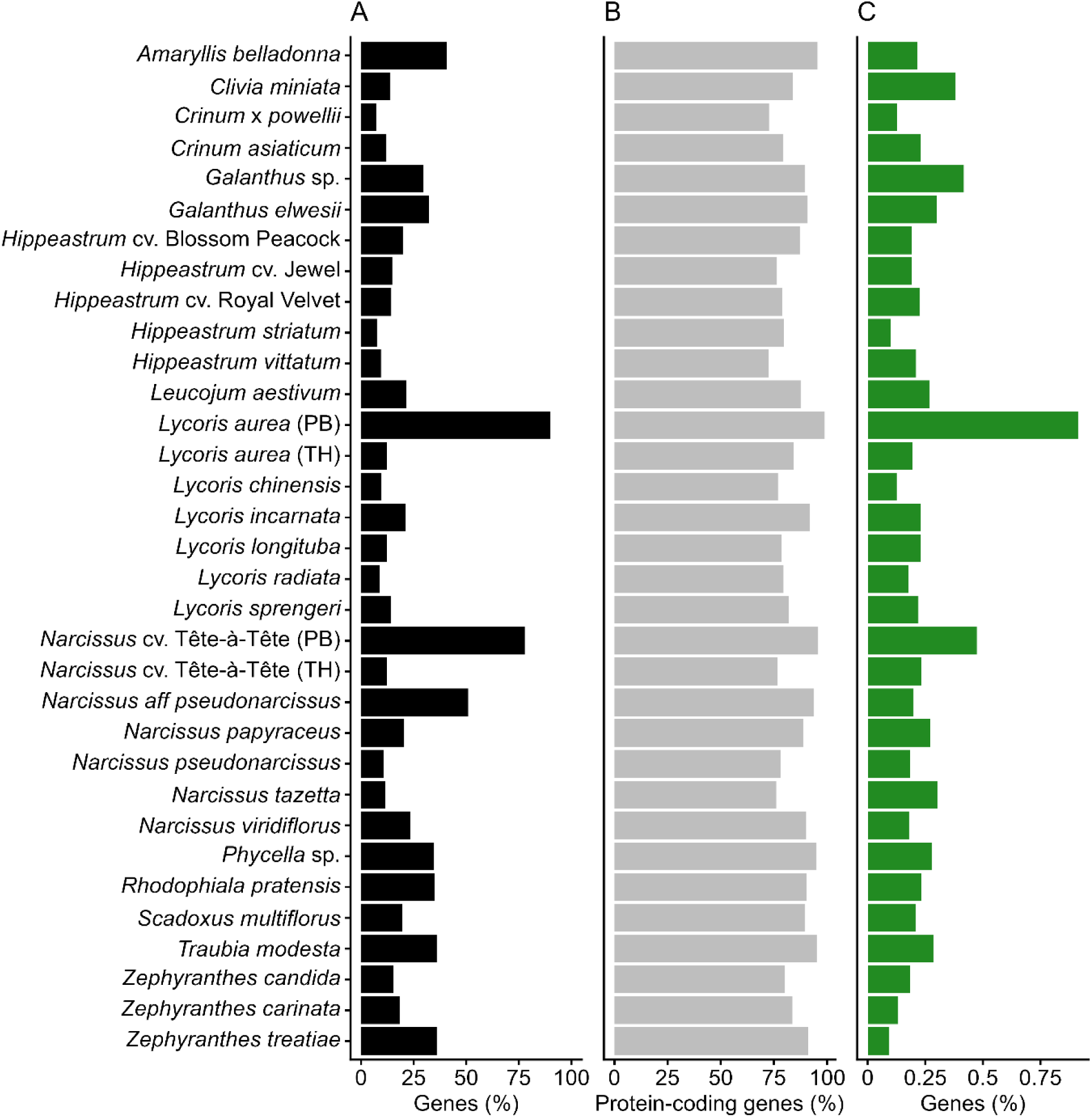
Proportion of “genes” annotated for each assembly. Proportion of (A) all trinity genes and (B) only protein coding genes annotated with protein-based datasets (EggNOG, Pfam-A and Uniprot SwissProt); If multiple peptides were predicted for a gene, the annotation of any of those peptide sequences was considered. (C) Proportion of trinity genes with at least one “isoform” annotated with Infernal using the Rfam dataset. For *Lycoris aurea* and *Narcissus* cv. ‘Tête-à-Tête’, two assemblies were generated, one using only long-read data (PB) and another using both long- and short read data with Trinity (TH).

### Technical validation

The assembly completeness was assessed with BUSCO 5.7.0 ^[103]^, using the liliopsida_odb10 dataset. This showed that the most complete assembly (single + duplicated) was that of *H*. cv. Royal Velvet (93.7%), followed closely by *Clivia miniata*, *Crinum* x *powellii*, *Z. candida*, *H.* cv. Jewel, *L. sprengeri* and *L. longituba* (Figure 5). The assemblies of *L. aurea* showed different proportions of complete BUSCOs, with the hybrid assembly having a proportion of complete BUSCOs higher by 15 points. Meanwhile, the assemblies of *N.* cv. ‘Tête-à-Tête’ showed similar completeness, the hybrid assembly was only 0.15% more complete than the long-read one. In the case of *L. aurea*, the hybrid assembly contained samples from multiple studies not pooled for long-read sequencing, while *N.* cv. ‘Tête-à-Tête’ datasets were all originated from the same sample and the mRNA used for the long-read sequencing was a pool of the different tissues of the plant. Conversely, three assemblies generated with data from the 1kP dataset lacked over 60% of the expected BUSCOs: *Traubia modesta* (71.1%), *A. belladonna* (66.6%), and *Z. treatiae* (65.6). Comparison to the original assemblies published by the OneKP iniative ^[67,105]^ showed that both the original and our re-assembled versions are similarly incomplete (Figure 6).

**Figure 6.**
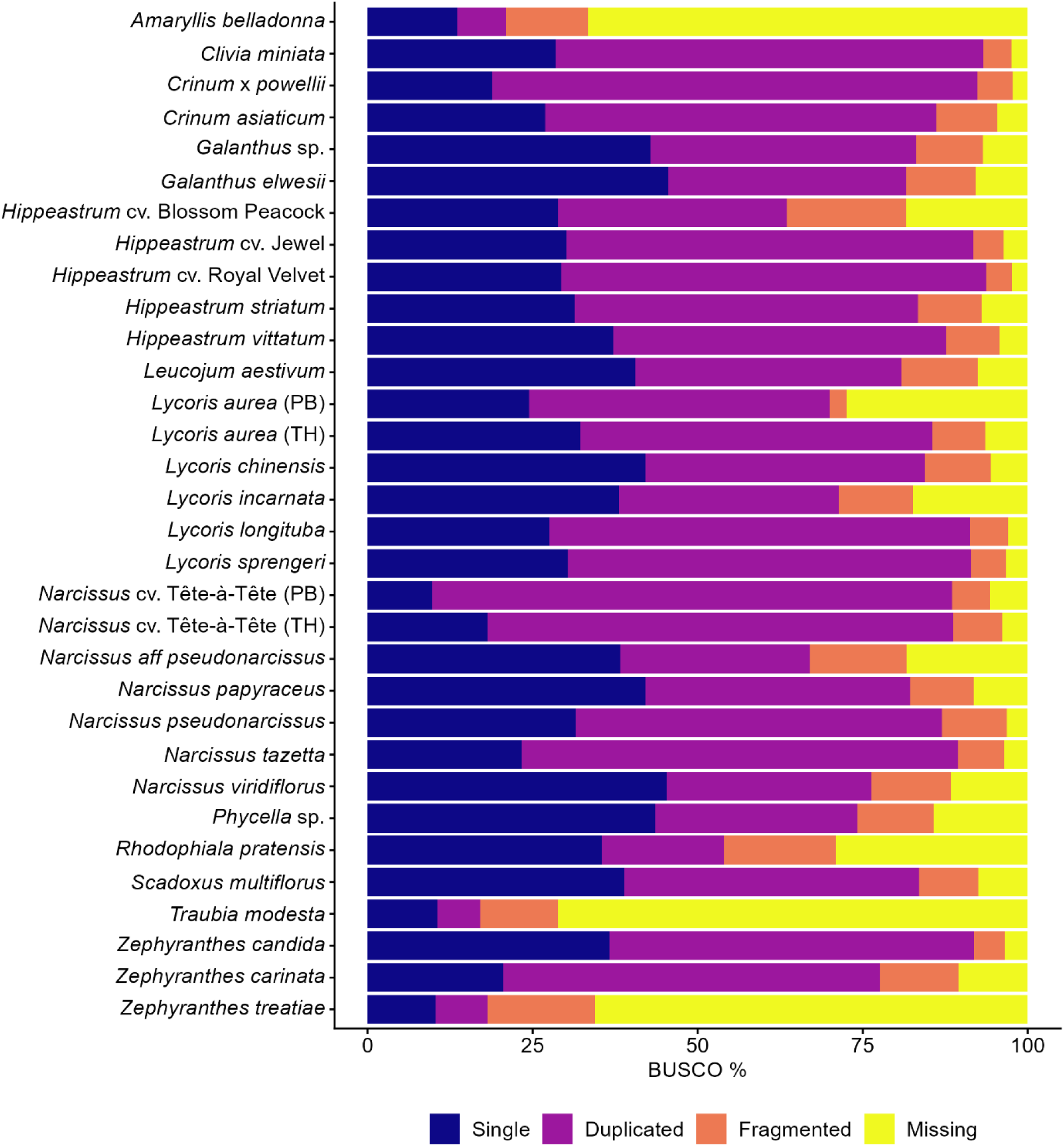
Assembly completeness evaluated with Benchmarking Universal Single-Copy Orthologs (BUSCO), using the liliopsida_odb10 dataset. For *Lycoris aurea* and *Narcissus* cv. ‘Tête-à-Tête’, two assemblies were generated, one using only long-read data (PB) and another using both long- and short read data with Trinity (TH).

Additionally, transcriptomes were analyzed with DOGMA 1.0.2 (DOmain-based General Measure for transcriptome and proteome quality Assessment, reference set for monocots), based on the analyses from RADIANT 1.1.5 with the dataset Pfam37.3. Of the 33 assemblies, 24 had scores greater than 80% (Figure 7) and 6 others showed completeness greater than 60%. As seen with BUSCO assessment, *A. belladonna*, *T. modesta* and *Z. treatiae* are the most incomplete assemblies, with scores of 24.9, 25.9 and 33.6%, respectively. The two organisms with both long and short read data had contrasting results: while for *L. aurea* the hybrid assembly had a score 10.1 higher compared to the long-read assembly, *N.* cv. ‘Tête-à-Tête’ long-read assembly had a greater (7.7 higher) completeness. Among the most complete, four assemblies had similar scores: *H.* cv. ‘Royal Velvet’*, C. miniata*, *Z. candida*, and *N.* cv. ‘Tête-à-Tête’ (PB). The low completeness of the transcriptomes from the 1kP dataset may be due to the effort required to obtain transcriptomes of many species simultaneously, which resulted in limiting sampling depth or tissues representation. However, this problem is not observed in other assemblies generated with a single replicate, such as *H. striatum*, *N. papyraceus* and *Scadoxus multiflorus*.

**Figure 7.**
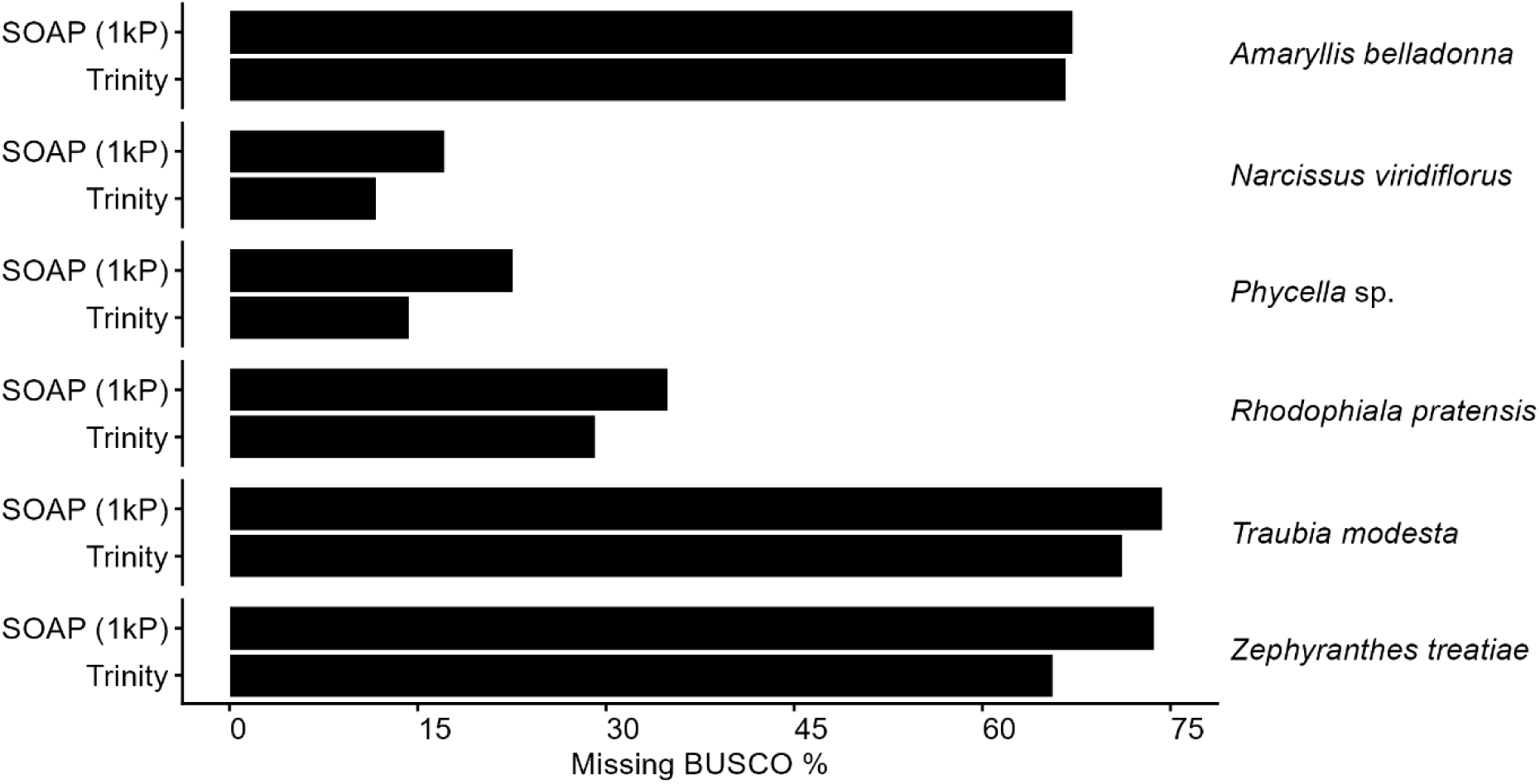
Little improvement was achieved by re-assembling data from the OneKP dataset^[67,105]^. Final assemblies of Amaryllidoideae species from the OneKP initiative were compared with de novo assemblies generated here, using the BUSCO dataset liliopsida_odb10.

The percentage of reads that map back to the assembly is another estimation of assembly quality is. Here, expression was estimated using two techniques: quasi-mapping with Salmon and pseudo-alignment with Kallisto (Figure 8). For all assemblies, mean mapping rate was higher with Salmon, reaching a maximum of 94.5% for *H.* cv. ‘Jewel’ and a minimum of 66.8% for *A. belladonna*. In the case of Kallisto, the highest mapping rate achieved was for *Phycella* sp. with 92.1%. Although the lowest Kallisto mean mapping rate was that of *Narcissus* cv. ‘Tête-à-Tête’ (TH), with a mean mapping rate of 40.9%, this result was the one that most differed between Kallisto and Salmon, with which the mapping rate was of 90.3%.

**Figure 8.**
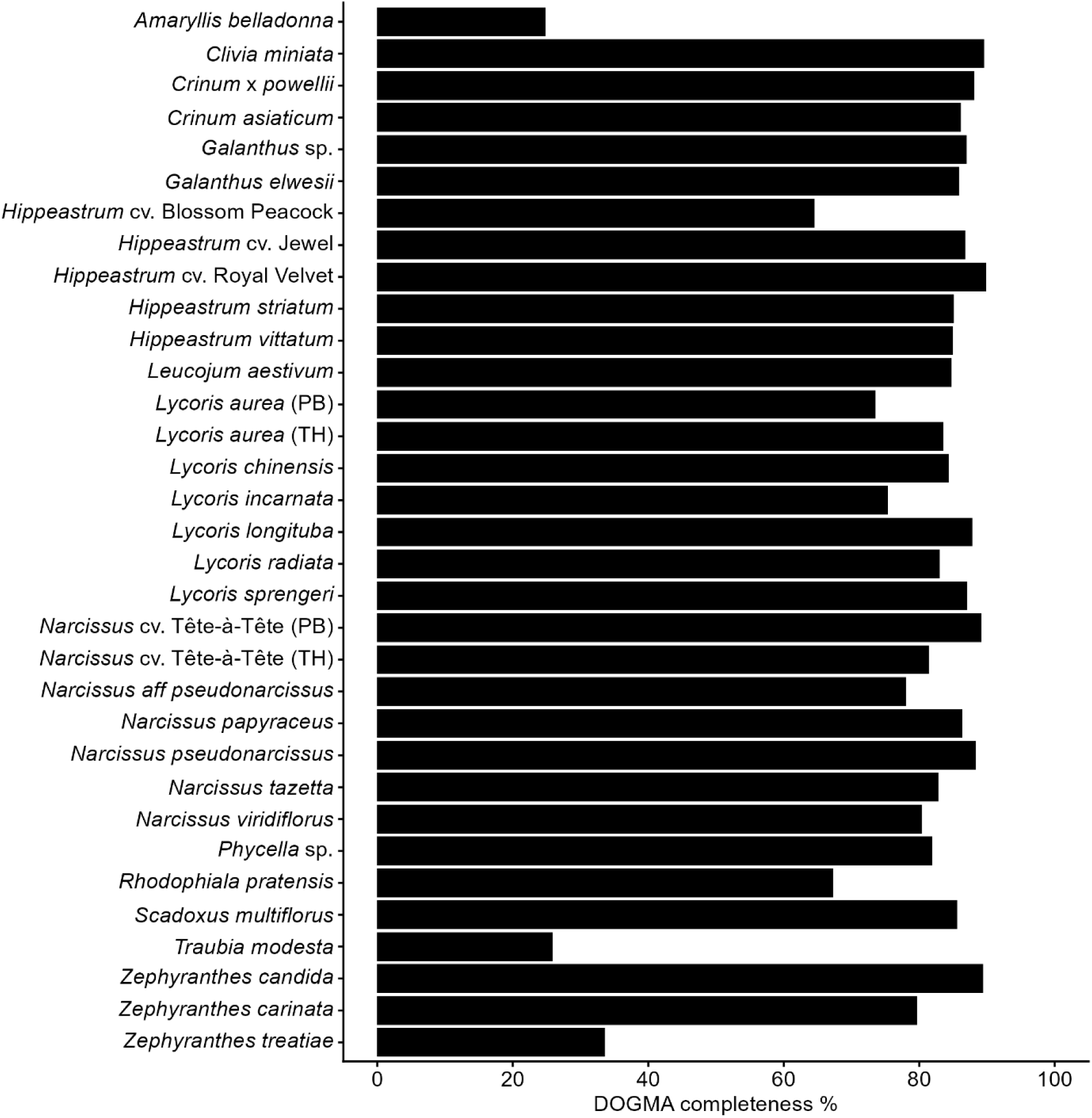
Most assemblies have moderate DOGMA completeness scores. For *Lycoris aurea* and *Narcissus* cv. ‘Tête-à-Tête’, two assemblies were generated, one using only long-read data (PB) and another using both long-and short read data with Trinity (TH). Data was assessed with RADIANT dataset for Pfam37.3 and the monocots reference set.

### Usage notes

This dataset can be used for mining genes of interest in one or many species of this family as well as for comparative transcriptomics. Fasta files for one or more species can be downloaded and used as a custom database for BLAST or other sequence-based analysis. This approach was employed in two previous publications of our team with the first version of this dataset to identify putative transcripts encoding for Norbelladine Synthase, Norcraugsodine/Noroxomaritidine Reductase^[119]^, Norbelladine 4’*O*-methyltransferase, and CYP96T (noroxomaritidine synthase)^[17]^ and Coclaurine *N*-methyltransferase-like (C*N*MT)^[120]^.

For each assembly, annotation results (based on BLAST against Uniprot, hmmscan against Pfam-A, emapper against eggNOG 5.0, infernal against Rfam 15.0; as well as predictions of transmembrane domains and signal peptides from TmHMM2 and SignalP6, respectively) can be extracted individually or from the annotation report, which includes all annotations, as well as gene ontology and KEGG pathway assignments based on the matches against Uniprot, Pfam-A and eggNOG. Since these are text files, users may perform text searches to find candidate genes by using dataset IDs, gene symbols or protein names.

Finally, expression quantification was conducted for each assembly and is provided as both read count and transcripts per million for both trinity “genes” and “isoforms” (except for both long read assemblies, in which case only “gene” expression is provided) estimated with Kallisto and Salmon. It is important to highlight that several factors impact the expression levels estimated, including biological, geographic, and technical. The data we provide has no batch effect correction. Users may perform batch effect correction or analyze the studies separately (by selecting the columns corresponding to the study of interest based on the SRA run ID, which are column names in these files).

To exemplify the usefulness of this dataset, we created a blast database containing the predicted protein sequences for all 33 assemblies and searched for proteins homologous to those involved in the Amaryllidaceae alkaloid pathway. Enzymes upstream of norbelladine synthase (NBS) were found in Uniprot SwissProt and used as baits for BLASTp. Given that several enzymes downstream of NBS have been recently characterized^[16–18,77,120]^, their sequences are not currently available in UniProt. Thus, their mRNA sequences, obtained from either Genbank or individual studies, were used as baits for BLASTx. These sequences and, when available, their accession numbers, are provided in Supplementary Data S1 and S2. Results were filtered to only keep matches with e-value < 1^-10^, sequence identity > 50%, and query coverage > 70%. Next, only the “best” protein sequence from each trinity “gene”, based on sequence completeness, length, and prediction score was selected.

Transcripts with homology to 19 enzymes known to catalyze 20 steps of the pathway (Figure 9) were found in 16 assemblies (namely, *C. miniata*, *C. asiaticum*, both *Galanthus* spp., *L. aestivum*, *Lycoris aurea* (PB), *L. incarnata*, *L. longituba*, *L. sprengeri*, *N. papyraceus*, *N. aff. pseudonarcissus*, *N. pseudonarcissus*, *N.* cv. ‘Tête-à-Tête’ (PB), and *S. multiflorus*), and in 10 assemblies candidate transcripts for 19 steps of the pathway were found. In *A. belladonna*, no acceptable matches (sequence identity > 50%, and query coverage > 70%) were found for seven enzymatic steps, including Phenylalanine Ammonia-Lyase and Tyramine Decarboxylase. Although two predicted peptides from this assembly had high sequence identity to Phenylalanine Ammonia-Lyase and one to Tyramine Decarboxylase, they were short and partial (missing either start or stop codon, or both). Additionally, the assemblies of *Z. treatiae* and *T. modesta* had no acceptable matches for 4 steps. These results agree with BUSCO and DOGMA scores, which showed these three were the least complete assemblies. Nevertheless, the partial matches found can be used to amplify the entire transcript, as was recently done for CNMT^[120]^.

**Figure 9.**
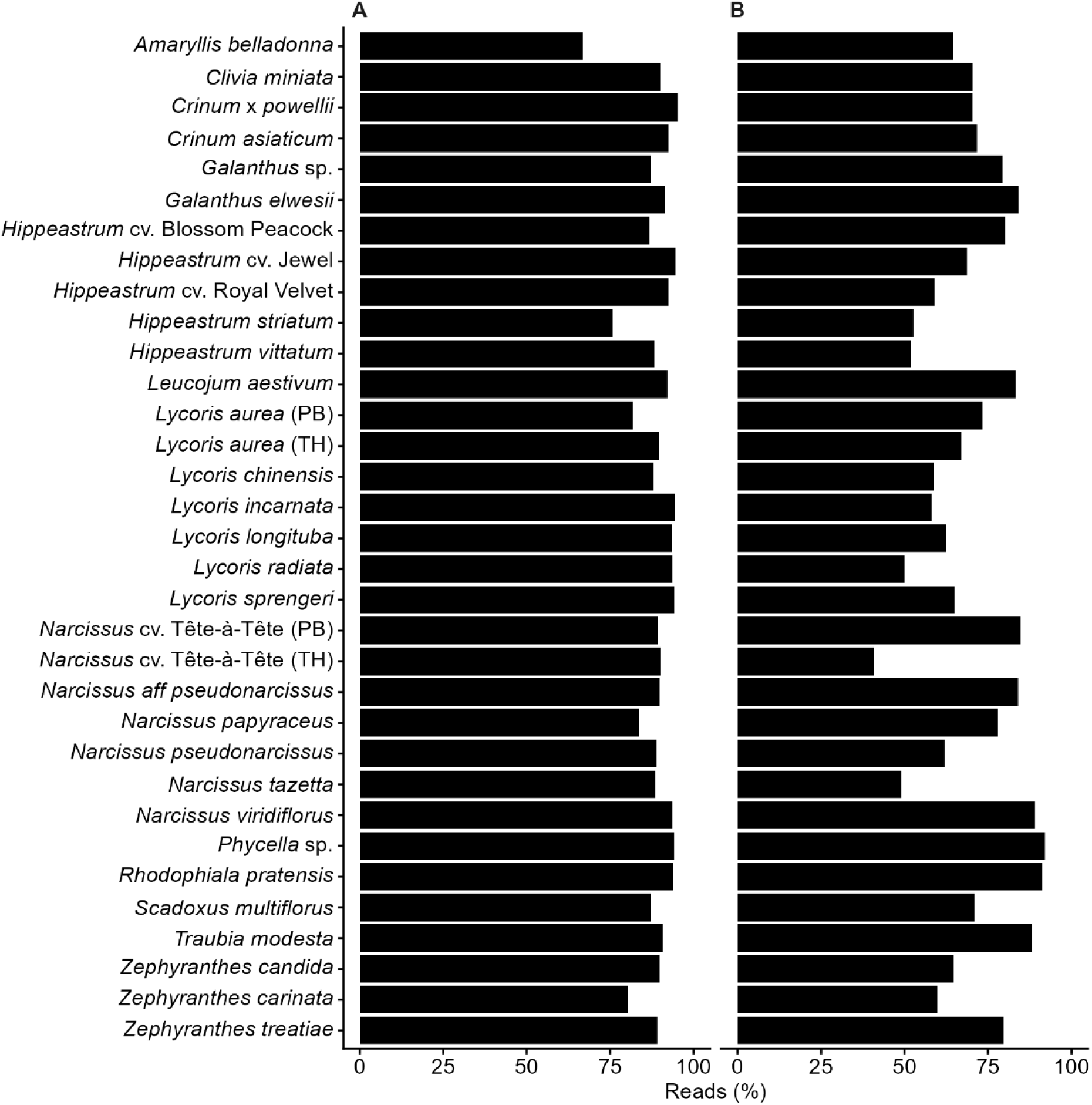
Most assemblies showed high rates of quasi-mapping (A) and moderate rates of pseudo-alignment (B). Bars indicate mean mapping percentage, dots indicate individual samples. For *Lycoris aurea* and *Narcissus* cv. ‘Tête-à-Tête’, two assemblies were generated, one using only long-read data (PB) and another using both long- and short read data with Trinity (TH).

**Figure 10.**
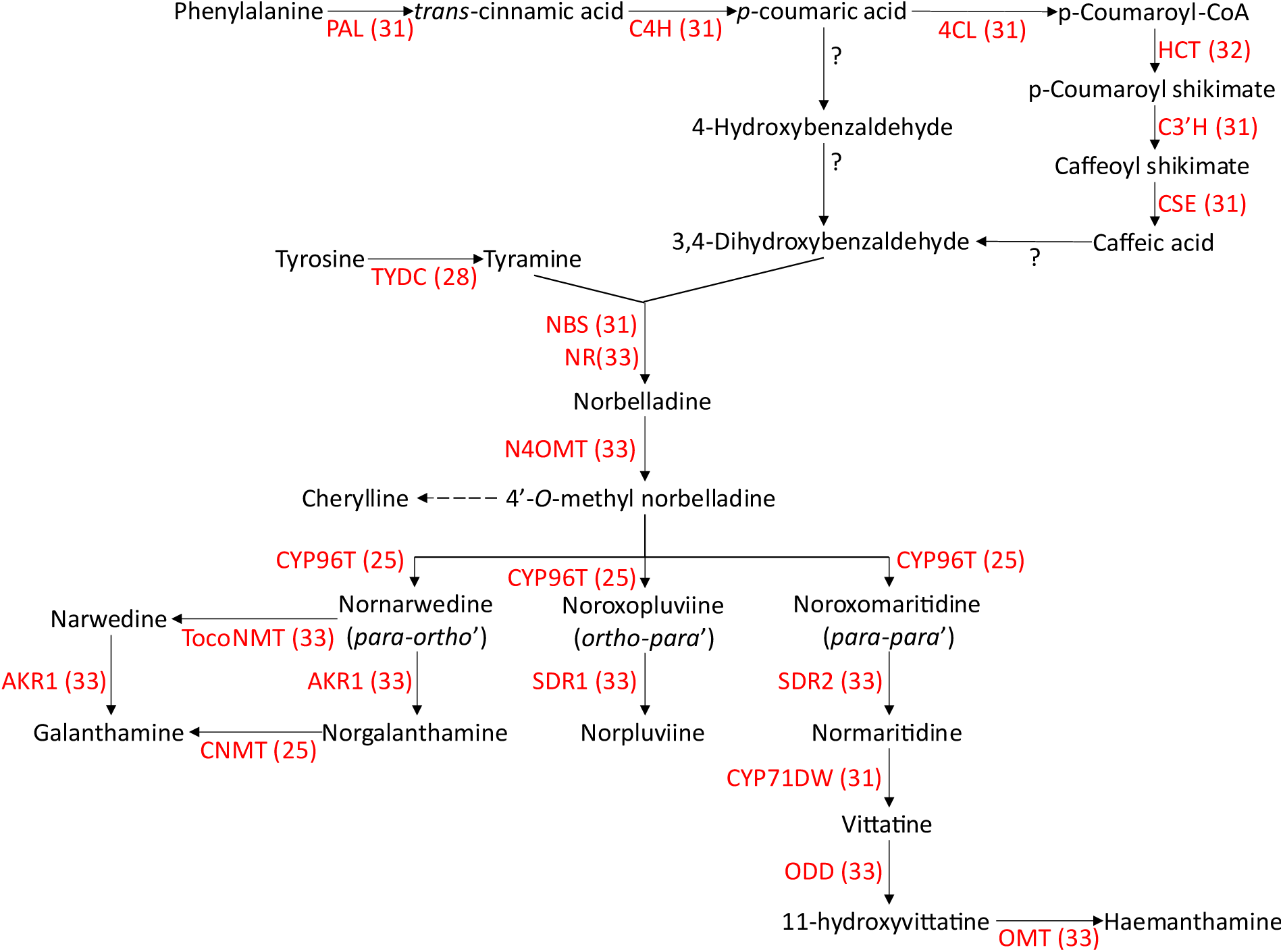
Predicted proteins putatively involved in the Amaryllidaceae alkaloid pathway were found in most assemblies of the AmarylOmicBase dataset. Metabolites are shown in black and enzyme acronyms in red. Number of assemblies in which candidate transcripts were found through BLASTP or BLASTX are shown inside parenthesis in red. Dashed line indicates multiple steps involved in the conversion of 4’-*O*-methyl norbelladine into cherylline. PAL: phenylalanine ammonia-lyase; C4H: cinnamate 4-hydroxylase; 4CL: 4-coumarate-CoA ligase; HCT: Hydroxycinnamoyltransferase; C3’H: *p-*coumaroyl 3′-hydroxylase; CSE: caffeoylshikimate esterase; TYDC: tyrosine decarboxylase; NBS: norbelladine synthase; NR: norcraugsodine/noroxomaritidine reductase; N4OMT: norbelladine 4’-*O*-methyltransferase; CYP96T: cytochrome P450 96T; Toco*N*MT: tocopherol *N*-methyltransferase; AKR1: aldo-keto reductase 1; *C*NMT: coclaurine *N*-methyltransferase-like enzyme; SDR1/2: short-chain alcohol dehydrogenase/reductase 1/2; CYP71DW: cytochrome P450 71DW; ODD: 2-oxoglutarate-dependent dioxygenase; *O*MT: *O*-methyltransferase; ?: unknown enzymes. Adapted from Liyanage, et al. ^[121]^.

One limitation of this dataset is the low completeness of several assemblies, namely *A. belladonna*, *T. modesta*, and *Z. treatiae*. Given that the assemblies generated for these species by One Thousand Plant Transcriptomes Initiative ^[67]^ had similar completeness scores, it is unlikely that the problem originates with the assembly pipeline. Although these transcriptomes are currently the only ones available for the *Amaryllis* and the *Traubia* genera and for *Z. treatiae*, users should be cautious when interpreting the failure of detection of a gene of interest.

Although we provide two assemblies for *N*. cv. ‘Tête-à-Tête’ and *L. aurea*, users should prioritize the highest quality assembly in each case. On one hand, the long-read assembly of *N*. cv. ‘Tête-à-Tête’ shows slightly better assessment results and while no acceptable match to C4H was found in the TH assembly. On the other hand, 27.4% of BUSCOs from the Liliopsida database were missing from the long-read assembly of *L. aurea*, compared to only 8% in the hybrid assembly. Regarding DOGMA, the TH assembly showed a completeness of 83.6%, a score 10.1% higher than the one observed for the long-read assembly. Although the TH assembly had a higher mapping rate with Salmon, the PB assembly performed better with Kallisto. Finally, given that no acceptable match to Norbelladine Synthase was found in PB assembly’s predicted proteome predicted, we suggest that users prioritize the hybrid assembly for *L. aurea*.

## Conclusion

Currently, the AmarylOmicBase consolidates 39 studies spanning 27 species, as well as four hybrid cultivars, into standardized and searchable resource, with assemblies, annotations, and expression quantification matrices, accompanied by reproducible scripts. By harmonizing disparate datasets, it enables rapid fishing of pathway candidates, cross-species comparison of isoenzymes, and targeted validation. For example, our proof-of-concept mining recovered transcripts encoding enzymes that catalyze 20 reactions in the Amaryllidaceae alkaloid pathway. Using conservative thresholds to filter blast results, we identified transcripts with homology to all known enzymes involved in the biosynthesis of these alkaloids in 16 assemblies included in the dataset, 10 other assemblies were missing candidate transcripts for only one enzymatic step. The database also exposes gaps due to sampling, sequencing depth, or assembly choices, guiding where new experiments or resequencing will have the highest payoff. Beyond alkaloids, the resource supports broader questions in specialized metabolism, from precursor supply to tailoring reactions and regulatory nodes. We encourage researchers on Amaryllidoideae transcriptomics to contact us when new data becomes available or to follow our pipeline to generate transcriptome assemblies and annotations that could then be included in an update of the AmarylOmicBase dataset.

## Data availability

Assemblies, predicted protein sequences, expression quantification, annotation results and the annotation compiled reports are available in Zenodo with the identifier https://doi.org/10.5281/zenodo.20349158 ^[115]^. Additionally, assemblies were deposited on NCBI Transcriptome Shotgun Assembly (when all raw data used for the assembly was also deposited in NCBI SRA) with the accession numbers^[122–147]^ indicated in Table 3. Expression data was deposited on NCBI GEO with the accession numbers GSE329951^[148]^ for *C.* × *powellii*, GSE329957^[149]^ for *L. aestivum*, GSE330014^[150]^ for *N. papyraceus*, and GSE331457^[151]^ for the remaining assemblies. Assemblies generated with data published in ENA EMBL (OneKP project) or in multiple databases (*L. radiata*, deposited on NCBI SRA and on NGDC GSA), were deposited with the expression data on NCBI GEO Super Series GSE331457^[151]^.

**Table 3.** *De novo* transcriptome assemblies generated in this study were deposited in NCBI TSA, when the raw data were originally deposited in NCBI SRA.

| Assembly name | TSA accession | Assembly name | TSA accession |
| --- | --- | --- | --- |
| *Clivia miniata* | DBNKRK000000000<sup>[122]</sup> | *Lycoris chinensis* | DBNKRE000000000<sup>[130]</sup> |
| *Crinum asiaticum* | DBNIJL000000000<sup>[146]</sup> | *L. incarnata* | DBNKRD000000000<sup>[130]</sup> |
| *C. × powellii* | DBNKRS000000000<sup>[123]</sup> | *L. longituba* | DBNKRG000000000<sup>[133]</sup> |
| *Galanthus* sp. | DBNKRR000000000<sup>[138]</sup> | *L. sprengeri* | DBNKRL000000000<sup>[135]</sup> |
| *G. elwesii* | DBNKRO000000000<sup>[136]</sup> | *Narcissus* aff. *pseudonarcissus* | DBNKRP000000000<sup>[137]</sup> |
| *Hippeastrum* cv. 'Blossom Peacock' | DBNMYE000000000<sup>[140]</sup> | *N. papyraceus* | DBNKRN000000000<sup>[147]</sup> |
| *H.* cv. 'Jewel' | DBNMYC000000000<sup>[142]</sup> | *N. pseudonarcissus* | DBNIJK000000000<sup>[129]</sup> |
| *H.* cv. 'Royal Velvet' | DBNMYD000000000<sup>[141]</sup> | *N.* cv. Tête-à-Tête (PB) | DBNNFO000000000<sup>[143]</sup> |
| *H. striatum* | DBNIJG000000000<sup>[125]</sup> | *N.* cv. Tête-à-Tête (TH) | DBNUFN000000000<sup>[144]</sup> |
| *H. vittatum* | DBNKRF000000000<sup>[132]</sup> | *Scadoxus multiflorus* | DBNIJH000000000<sup>[126]</sup> |
| *Leucojum aestivum* | DBNIJJ000000000<sup>[128]</sup> | *Zephyranthes candida* | DBNKRH000000000<sup>[134]</sup> |
| *Lycoris aurea* (PB) | DBNFTY000000000<sup>[124]</sup> | *Z. carinata* | DBNIJI000000000<sup>[127]</sup> |
| *L. aurea* (TH) | DBNKRI000000000<sup>[139]</sup> |  |  |

## Code availability

Scripts for read quality control, assembly, annotation, and report are available on the Gitlab repository KarenGoncalves/AmarylOmicBase^[152]^.

## Author Contributions Statement

Conceptualization: KCGdS, NM, IDP; Methodology: KCGdS; Validation: KCGdS, NM; Formal analysis: KCGdS; Investigation: KCGdS, NM; Resources: IDP; Data curation: KCGdS; Writing – Original Draft: KCGdS; Writing – Review & Editing: KCGdS, NM, IDP; Visualization: KCGdS, NM; Supervision: IDP; Project administration: NM, IDP; Funding acquisition: IDP.

## Acknowledgements

The authors gratefully acknowledge the research groups and consortia who generated and made publicly available the RNA sequencing datasets used in this study. Their contributions to open data sharing have made this work possible. This research was enabled in part by support provided by the Digital Research Alliance of Canada (alliancecan.ca), through access to advanced research computing resources and technical support. Thanks are extended to the Canadian taxpayers and the Canadian government for supporting the Canada Research Chairs Program.

